# The kinetics of bedaquiline diffusion in tuberculous cavities opens a window for emergence of resistance

**DOI:** 10.1101/2024.10.09.617456

**Authors:** Annamarie E. Bustion, Jacqueline P. Ernest, Firat Kaya, Connie Silva, Jansy Sarathy, Landry Blanc, Marjorie Imperial, Martin Gengenbacher, Min Xie, Matthew Zimmerman, Gregory T. Robertson, Danielle Weiner, Laura E. Via, Clifton E. Barry, Radojka M. Savic, Véronique Dartois

## Abstract

Cavitary tuberculosis (TB) is difficult to cure and a site of relapse. Bedaquiline has been a wonder drug in the treatment of multidrug resistant TB, but emergence of resistance threatens its sustained success. To investigate the role of drug distribution in resistance development, we designed a novel laser-capture microdissection scheme to spatially resolve the penetration of bedaquiline in the necrotic center (*caseum*) of cavities, a recalcitrant site of disease. Working with preclinical models that form large necrotic lesions, we profiled bedaquiline and two next generation diarylquinolines TBAJ-587 and TBAJ-876. Drug concentrations were measured in laser-captured areas of cavity caseum as a function of time and distance from blood supply. To simulate drug coverage in patient cavities, the data were modeled, and drug penetration parameter estimates were linked to clinical plasma pharmacokinetics for bedaquiline and the new diarylquinolines. Pharmacokinetic-pharmacodynamic (PK-PD) simulations revealed that bedaquiline reaches efficacious concentrations in outer and deep caseum after several weeks to months and lingers at subtherapeutic concentrations up to 3 years after therapy ends. TBAJ-587 and TBAJ-876, currently in clinical development, achieve bactericidal concentrations in caseum more rapidly and shorten the window of suboptimal concentrations post treatment compared to bedaquiline. Simulations of clinically plausible dosing schemes were conducted to guide the design of clinical trials for cavitary TB and help mitigate resistance development. In summary, the slow kinetics of diffusion of bedaquiline into and out of cavity caseum creates spatio-temporal windows of subtherapeutic concentrations. Site-of-disease simulations of TBAJ-587 and TBAJ-876 predict reduced opportunities for resistance development.

**SIGNIFICANCE:** Clinical resistance to bedaquiline has emerged faster than anticipated. Understanding potential contributing factors could help curb further resistance development, not only for bedaquiline but also for the next generation diarylquinolines currently in phase 2, TBAJ-587 and TBAJ-876. Here we hypothesized and confirmed that the extended time to reach steady state and slow clearance of bedaquiline leads to extremely slow diffusion into and out of cavity caseum, a recalcitrant site of TB disease and relapse. Through modeling of experimental data in a preclinical model of cavitary TB and clinical simulations, we show that the next generation diarylquinolines may reduce spatio-temporal windows of resistance development compared to bedaquiline. Our results can inform dosing schemes of diarylquinoline-based therapies that limit resistance development.

## INTRODUCTION

Cavitary tuberculosis (TB) disease is associated with unfavorable outcomes (1, 2). Profound drug tolerance of the resident bacterial population (3) and limited drug diffusion within caseum, a non-vascularized site-of-disease (4, 5), are considered key factors driving poor treatment response in cavitary patients (6). Central caseum is also found in closed nodular lesions or granulomas. In the pre-antibiotic era, autopsies following TB-related deaths revealed variable but generally high bacterial burden in the caseous foci of necrotic granulomas and cavities, distributed throughout the caseous mass (7-9). Thus, complete diffusion of bactericidal agents throughout the necrotic core of these recalcitrant lesion types appears critical to achieve durable cure. Indeed, cavity caseum was recently shown as the major site of relapse {Malherbe, 2024 #5254}.

Bedaquiline (BDQ) has been the wonder drug of MDR- and XDR-TB treatment, contributing to treatment shortening from 24 to 6 months by the BDQ-pretomanid-linezolid combination (in a trial known as NIX-TB) (10). It is the backbone of numerous combination trials (11-13) to identify regimens that further reduce treatment duration, with a few promising but preliminary results to be confirmed in larger trials and diverse patient populations (14, 15). Despite this, BDQ suffers from pharmacokinetic (PK) liabilities: its extreme hydrophobic and cationic amphiphile properties result in high binding to plasma proteins and tissue macromolecules (16), extensive tissue distribution, and long terminal half-life. Lingering, subtherapeutic concentrations have been measured in plasma and at the site of disease for several months after the end of therapy (17, 18), conditions that are prone to resistance acquisition. Indeed, emergence of BDQ resistance has occurred surprisingly quickly across patient populations (19, 20).

In previous work, we have shown that BDQ as well as clofazimine, a drug with similar physicochemical and PK properties, exhibit strong differential partitioning in cavity wall and cavity caseum, due to high accumulation in immune cells and poor diffusion through necrotic foci (5, 21-24). Given the clear positive impact of BDQ on treatment outcome but the disappointing results of the 14-day early bactericidal activity trial (25), we hypothesize that BDQ exhibits very slow kinetics of diffusion into caseum, concomitant with gradual redistribution from the cellular rim into the caseous core of cavities and large necrotic nodules, the ‘hard-to-treat’ disease sites.

The BDQ analogs and next generation diarylquinolines TBAJ-587 and TBAJ-876 have improved pharmacological profiles (26) and are in clinical development to replace BDQ and accelerate cure. They were designed to achieve lower lipophilicity, higher clearance, and similar or increased potency compared to BDQ. They are more potent against replicating bacteria and non-replicating persisters in caseum (27), and improve sterilization in the relapse mouse model when replacing BDQ (28).

To investigate the potential impact of diarylquinoline distribution on efficacy and emergence of resistance at the site of cavitary disease, we first resolved the spatio-temporal penetration of the three agents in the caseous areas of pulmonary lesions in a rabbit model of active TB. To quantify the kinetics of drug diffusion, we devised a laser-capture microdissection (LCM) strategy relying on distance maps generated by Euclidean Distance Transform. We then related achieved concentrations to the minimum bactericidal concentrations required to kill non-replicating persister bacteria in *ex vivo* caseum and applied the drug diffusion model to visualize target attainment in patient cavities. Simulating the caseum pharmacokinetic-pharmacodynamic (PK-PD) relations for the three diarylquinolines during the intensive and continuation treatment phase and post-therapy allowed for the comparison of temporal windows during which sub-therapeutic concentrations are present in caseum, creating opportunities for resistance development.

## RESULTS

### Slow kinetics of diarylquinoline diffusion into the necrotic centers of TB lesions

Autopsies from the pre-antibiotic era revealed large numbers of Mtb bacilli, observed as both single cells and clumps, throughout the necrotic center (or caseum) of closed nodules and cavities. Similar observations were made in rabbits with cavitary TB (**Figure S1**). We also know that non-replicating Mtb bacilli residing in caseum are profoundly tolerant to most antibiotic classes (27, 29). Thus, complete drug diffusion throughout non-vascularized necrotic foci at concentrations adequate to kill the resident bacteria is important to deliver cure.

In previous studies of BDQ penetration into large necrotic lesions of C3HeB/FeJ mice, we observed steep decreasing concentration gradients from the cellular rim into the necrotic center of caseous type I granulomas upon short term exposure (22, 30). Given the time required for BDQ to reach steady state in plasma, and its low free fraction, we hypothesized that it may exhibit slow diffusion kinetics into non-vascularized caseum and that the rabbit model of cavitary TB would be an adequate tool to resolve the spatio-temporal kinetics of diffusion.

In MDR-TB patients, BDQ is dosed at 400 mg daily for 14 days followed by 200 mg thrice weekly for 22 weeks. To identify the rabbit dose and the minimum number of doses that would reproduce human exposure at steady state, a series of dose finding PK studies were performed in naïve rabbits at 20, 80 and 160 mg/kg for 1 to 17 days (**Table S1**). Through modeling of the PK data and multi-dose simulations, we determined that 3 daily flat doses of 400 mg/rabbit achieved the C_ss[0-24]_ (average steady state concentration over the 24h dosing interval) target of 1000 ng/mL for BDQ. Long term dosing of 14 days or longer at 20 mg/kg similarly reproduced human exposure at steady state. For TBAJ-587 and TBAJ-876, the target AUCs were of 15 and 4 μg*h/mL, respectively, based on efficacy data in the mouse model of chronic TB (31, 32). Modeling of the plasma PK data (**Figure S2**) and multidose simulations predicted that the target AUCs would be achieved after 3 daily doses of 125 mg/kg TBAJ-587 and 100 mg/kg TBAJ-876. A follow up PK study in uninfected rabbits revealed more than dose-proportional exposure, from which 3 daily flat doses of 300 mg for TBAJ-587 and 125 mg for TBAJ-876 were recommended to achieve the desired exposure.

Next, cohorts of New Zealand White rabbits were infected via the aerosol route with Mtb strain HN878 as described (33), and disease was allowed to progress for 12 to 40 weeks, until typical lung lesions were present including large caseous lesions and cavities. At that time, rabbits received 1 to 28 doses of BDQ, TBAJ-587 or TBAJ-876. For animals who received 13 or more doses, the daily dose was reduced to 1/10 of the target dose (**Table S1**) to prevent excessive healing and resorption of caseous foci. This was accounted for in data analysis and modeling. To quantify the concentration gradients of these drugs in caseum as a function of time and space, large necrotic lesions were collected from infected rabbits at various time points after the last dose and subjected to laser-capture microdissection (LCM), to sample concentric areas of caseum equidistant from the interface with the cellular rim (**Figure 1A**). Each diarylquinoline and its active metabolite were quantified in LCM samples by LC/MS-MS, providing absolute concentrations as a function of distance from the edge of the necrotic region, where blood supply ends (**Figure 1B**). We observed steep concentration gradients after 1 and 3 doses, decreasing from the interface with the vascularized cellular rim towards deep caseum layers, which gradually became less steep as the number of doses increased to 14 and 28, for both parent and active metabolites. The gradient steepness also depended on distance from the caseum edge (**Figure 1B**).

**Figure 1.**
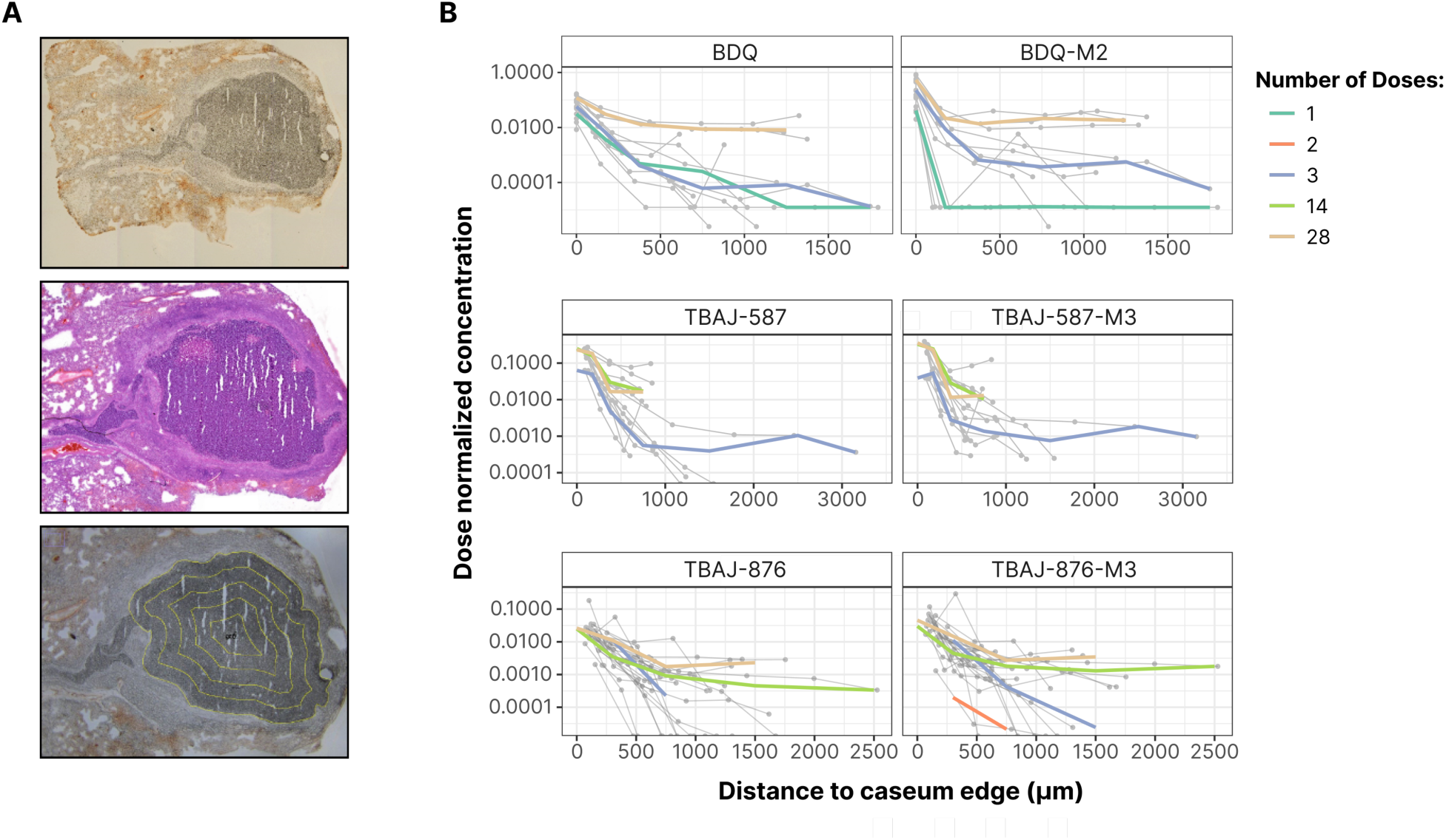
Diarylquinoline penetration in the caseous center of necrotic rabbit lesions. **(A)** Laser-capture microdissection (LCM) scheme to quantify the study drugs in caseum as a function of distance from the vascularized border. Image J Distance Map function was used to draw equidistant concentric areas to guide LCM sampling and calculate the surface of each sampled area. The average distance from the outer border of the caseum (in μm, X axis) was calculated by creating a caseum ‘mask’ and processing it using the Exact Euclidean Distance Transform plugin of the ImageJ software. **(B)** Concentrations of bedaquiline (BDQ), TBAJ-587, TBAJ-876, and their active metabolites in caseous foci as a function of distance from the interface between the caseous mass and the cellular (vascularized) rim. Experimental observations from individual rabbits are shown as dots and grey lines, and the median of raw data stratified by the total number of doses are represented as colored lines.

To determine whether a similar evolution of concentration gradient is recapitulated in other animal models with large necrotic lesions and cavities, we dosed TB-infected C3HeB/FeJ mice and common marmosets with BDQ at the human equivalent doses (25 and 20 mg/kg, respectively) for 1 and 17 days (mice) and 31 and 36 days (marmosets). As seen in rabbits, the steepness of the concentration gradient from the outer rim of the caseum into the core was inversely proportional to the number of doses (**Figure S3**) and indicated that several weeks of daily dosing are required for BDQ to reach steady state at the center of necrotic lesions. Thus, the observed trend was recapitulated in three disease models. From there on, the rabbit model was selected for in-depth investigations.

### Quantitative modeling of diarylquinoline diffusion into caseum

For the three parent drugs and their metabolites, the rabbit plasma PK was best described by an oral two-compartment distribution model. Depending on the drug, the absorption phase was best modeled with 1 or 2 transit compartments (**Table 1**). BDQ and TBAJ-587 exhibited linear elimination, while TBAJ-876 clearance was best described using Michaelis Menten kinetics. A metabolite compartment was included to describe the metabolite concentrations (**Figure 2A**). For both parent and metabolite, a mathematical relationship was established to describe the extent of drug partitioning into caseum relative to the distance from the caseum edge at the interface with the cellular rim (**Equations 1-2**).

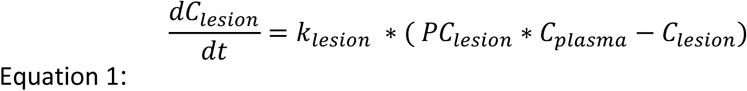

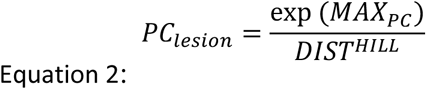

**Figure 2.**
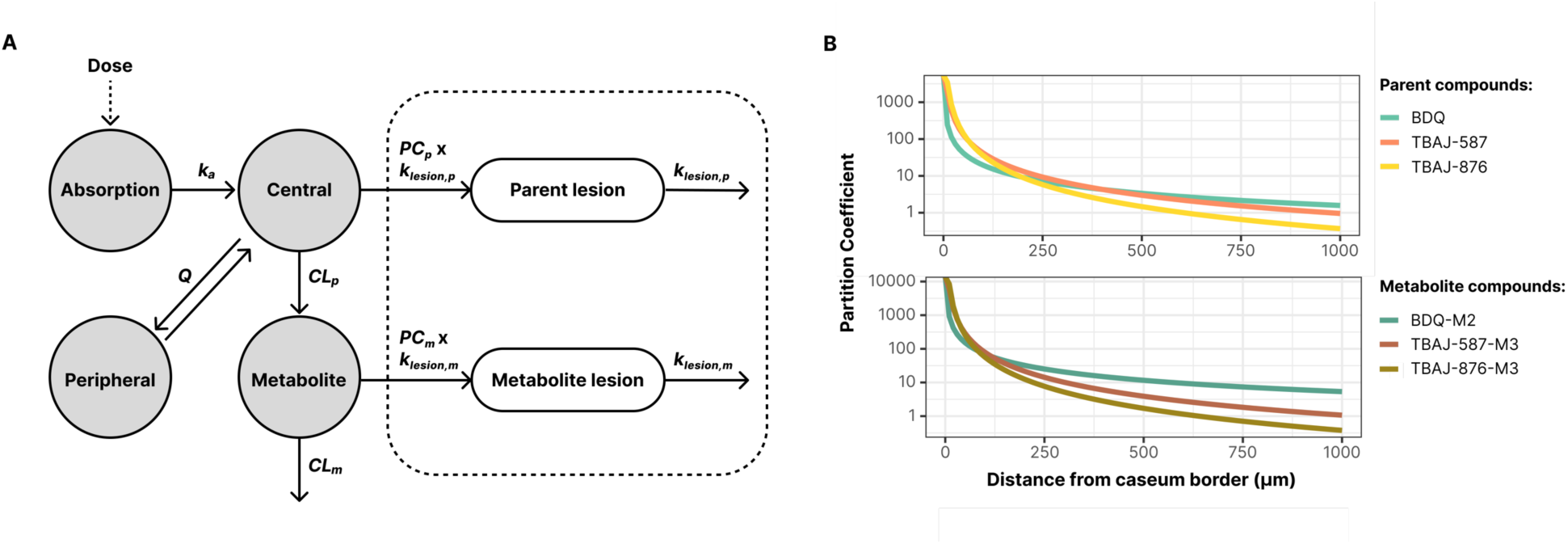
Diarylquinoline parent and metabolite plasma-to-lesion structural model. **(A)** Plasma PK for the diarylquinolines was described by an oral two-compartment distribution model. PK modeling of lesion compartments was implemented as an effect compartment to describe drug distribution from plasma (Equation 1). The final model included a distance covariate on the partition coefficient (Equation 2). **(B)** The relationship between partition coefficient and distance from the caseum border (Equation 2) was estimated for each diarylquinoline. TBAJ-587 and TBAJ-876 were found to have shallower gradients compared to BDQ.

**Table 1.** Rabbit plasma PK structural model and parameters.

| BDQ |  | TBAJ-587 |  | TBAJ-876 |  |
| --- | --- | --- | --- | --- | --- |
| 2 compartments, linear elimination, 1 transit compartment, metabolite (M2) compartment |  | 2 compartments, linear elimination, 1 transit compartment, metabolite (M3) central compartment and metabolite (M3) peripheral compartment |  | 2 compartments, saturated elimination, 2 transit compartments, metabolite (M3) compartment |  |
| PK parameter | Value (% RSE) | PK parameter | Value (% RSE) | PK parameter | Value (% RSE) |
| CL/F (L/h) | 11.19 (9.8) | CL/F (L/h) | 7.12 (13) | CL/F (L/h) | 37.26 (11) |
| - | - | - | - | K<sub>M</sub> (mg/L) | 0.195 (39) |
| V<sub>C</sub>/F (L) | 257.3 (21) | V<sub>C</sub>/F (L) | 89.4 (12) | V<sub>C</sub>/F (L) | 207.2 (18) |
| K<sub>TR</sub> (1/h) | 0.41 (18) | K<sub>TR</sub> (1/h) | 0.283 (18) | K<sub>A</sub> (1/h) | 1.39 (21) |
| - | - | - | - | MTT (h) | 1.43 (10) |
| Q/F (L/h) | 26.03 (62) | Q/F (L/h) | 13.46 (31) | Q/F (L/h) | 40.43 (10) |
| V<sub>P</sub>/F (L) | 301.5 (34) | V<sub>P</sub>/F (L) | 182.3 (30) | V<sub>P</sub>/F (L) | 1324 (22) |
| Metabolite CL/F (L/h) | 27.22 (11) | Metabolite CL/F (L/h) | 12.79 (12) | Metabolite CL/F (L/h) | 60.29 (12) |
| Metabolite V<sub>C</sub>/F (L) | fixed to VC | Metabolite V<sub>C</sub>/F (L) | 38.86 (54) | Metabolite V<sub>C</sub>/F (L) | 607.9 (12) |
| - | - | Metabolite V<sub>P</sub>/F (L) | 352.8 (20) | - | - |
| - | - | Metabolite Q/F (L/h) | 41.31 (25) | - | - |
| Parent Prop. Error (%CV) | 27.6 (7.8) | Parent Prop. Error (%CV) | 47.3 (6.5) | Parent Prop. Error (%CV) | 51.5 (10) |
| Metabolite Prop. Error (%CV) | 3.88 (16) | Metabolite Prop. Error (%CV) | 38.7 (10) | Metabolite Prop. Error (%CV) | 20.1 (7.9) |
| F IIV (%CV) | 31.8 (25) | F IIV (%CV) | 63.7 (22) | - | - |
| V<sub>P</sub>/F IIV (%CV) | 57.2 (16) | - | - | - | - |
| - | - | - | - | CL/F IIV (%CV) | 29.9 (15) |
| - | - | - | - | Metabolite CL/F IIV (%CV) | 35.1 (14) |

RSE, relative standard error; CL/F, apparent clearance; K<sub>M</sub>, Michaelis-Menten constant; V<sub>C</sub>/F, apparent volume of distribution of central compartment; K<sub>TR</sub>, absorption rate constant for transit compartment(s); K<sub>A</sub>, absorption rate constant; MTT, mean transit time; Q/F, apparent intercompartmental clearance; V<sub>P</sub>/F, apparent volume of distribution of peripheral compartment; Prop. Error, residual error model where the magnitude of error is proportional to predicted values; %CV, percentage of coefficient of variation; IIV, interindividual variability.

Plasma and lesion PK parameter estimates are summarized in **Tables 1** and **2**. Model development was guided by objective function value, goodness of fit plots, and VPCs (**Figures S4 and S5**). VPCs demonstrated the appropriateness of the final plasma and lesion models and fit to the raw data (**Figure S5**). The final models show that BDQ exhibits the steepest concentration gradients, while TBAJ-587 and TBAJ-876 follow a slightly shallower gradient at the caseum edge (**Figure 2B**), in line with their 10-fold higher free fraction (**Table 3**) and lower hydrophobicity compared to BDQ.

**Table 2.** Rabbit caseous lesion PK parameters.

| Parameter | BDQ<br>Estimate (RSE%) | TBAJ-587<br>Estimate (RSE%) | TBAJ-876<br>Estimate (RSE%) |
| --- | --- | --- | --- |
| $K_{\text{lesion,parent}}$ (1/h) | 0.001622 (148) | 0.006014 (34) | 0.3568 (10) |
| $\text{MAXPC}_{\text{parent}}$ | 8.011 (22) | 11.29 (6.3) | 12.73 (15) |
| $\text{HILL}_{\text{parent}}$ | 1.094 (12) | 1.641 (8.5) | 1.987 (18) |
| $K_{\text{lesion\_metabolite}}$ (1/h) | 0.003185 (79) | 0.01166 (27) | 0.3489 (7.1) |
| $\text{MAXPC}_{\text{metabolite}}$ | 9.387 (11) | 12.97 (6.3) | 14.05 (8.7) |
| $\text{HILL}_{\text{metabolite}}$ | 1.116 (10) | 1.868 (8.3) | 2.175 (11) |
| Proportional error<sub>parent</sub> (%CV) | 2.24 (31) | 24.42 (26) | NA |
| Add error<sub>parent</sub> (mg/L) | 1.31 (13) | 0.06 (74.8) | 0.91 (20) |
| Proportional error<sub>metabolite</sub> (%CV) | 5.28 (24) | 20.71 (24) | NA |
| Add error<sub>metabolite</sub> (mg/L) | 1.08 (16) | 0.06 (86) | 0.71 (28) |

RSE, relative standard error; $K_{\text{lesion}}$ , rate of partitioning from plasma into lesion; MAXPC, maximum extent of partitioning from plasma into lesion; HILL, exponent that determines the steepness of the curve that describes the extent of partitioning relative to distance from caseum edge; Prop. Error, proportional residual error model where the magnitude of error is *proportional* to predicted values; Add. Error, additive residual error model where the magnitude of error is constant across predicted values; %CV, percentage of coefficient of variation.

**Table 3.** Pharmacodynamic targets and plasma protein binding.

| | MIC <sup>[a]</sup> | casMBC<sub>90</sub><br>day 7 | casMBC<sub>90</sub><br>day 14 | casMBC<sub>50</sub><br>day 14 | casMBC<sub>10</sub><br>day 14 | Plasma $f_u$<br>(human, %) |
| --- | --- | --- | --- | --- | --- | --- |
| | $\mu\text{M}$ ($\mu\text{g/mL}$) | | | | | |
| BDQ | 0.18 (0.10) | 65 (36) | 4.14 (2.29) | 2.34 (1.30) | 1.32 (0.74) | 0.0053 |
| BDQ-M2 | 0.87 (0.47) | 38 (21) | 3.68 (1.99) | 1.50 (0.81) | 0.61 (0.33) | 0.035 |
| TBAJ-587 | 0.016<br>(0.01) | 1.5 (0.9) | 0.54 (0.33) | 0.45 (0.28) | 0.37 (0.23) | 0.034 |
| TBAJ-587-M3 | 0.18 (0.11) | 32 (19) | 2.49 (1.49) | 0.13 (0.08) | 0.007<br>(0.004) | 0.23 |
| TBAJ-876 | 0.015<br>(0.01) | 0.5 (0.3) | 0.15 (0.10) | 0.12 (0.08) | 0.10 (0.07) | 0.043 |
| TBAJ-876-M3 | 0.047<br>(0.03) | 2.0 (1.3) | 0.50 (0.32) | 0.46 (0.30) | 0.42 (0.27) | 0.4 |

<sup>[a]</sup> Microplate Alamar Blue Assay against H37Rv. $f_u$ : fraction unbound

### PK-PD simulations of diarylquinolines in caseous lesions during and post treatment

To relate concentrations achieved to concentrations required to reduce the bacterial burden in large necrotic lesions and cavities in TB patients, we first measured the bactericidal activity of BDQ, the TBAJs, and their active metabolites against non-replicating Mtb persisters in *ex vivo* caseum (3), after 7 and 14 days of incubation given the documented slow onset of BDQ’s bactericidal activity (34, 35) (**Figure S6**). Concentrations that reduce the bacterial load by 90, 50 and 10% (casMBC_90_, casMBC_50_ and casMBC_10_) were inferred from the raw data using a four-parameter logistic model for dose response (**Table 3, Figure S7**).

To estimate lesion concentrations in patients, the plasma-to-caseous lesion model parameters were linked to population PK models for BDQ (36), TBAJ-587 (37) and TBAJ-876 ((38) and present work). The standard BDQ dosing regimen of 400 mg once daily for 2 weeks followed by 200 mg thrice weekly for 22 weeks was simulated in 500 virtual patients, and 500 virtual patients received either 50, 100, or 200 mg TBAJ-587 or 25, 50, or 100 mg TBAJ-876 once daily for six months. Average concentration ranges in plasma and caseum were simulated during the buildup phase (**Figure 3A**) and concentration gradients as a function of distance from the cellular rim were simulated over a 6-month treatment period and up to 3.5 years after the end of therapy (**Figure 3B**). Average daily concentrations achieved at increasing depth in necrotic foci were related to the casMBC_90_, casMBC_50_, and casMBC_10_, and revealed distinct patterns of target attainment relative to distance and time for the three drugs and their active metabolites (**Figure 3C**).

**Figure 3.**
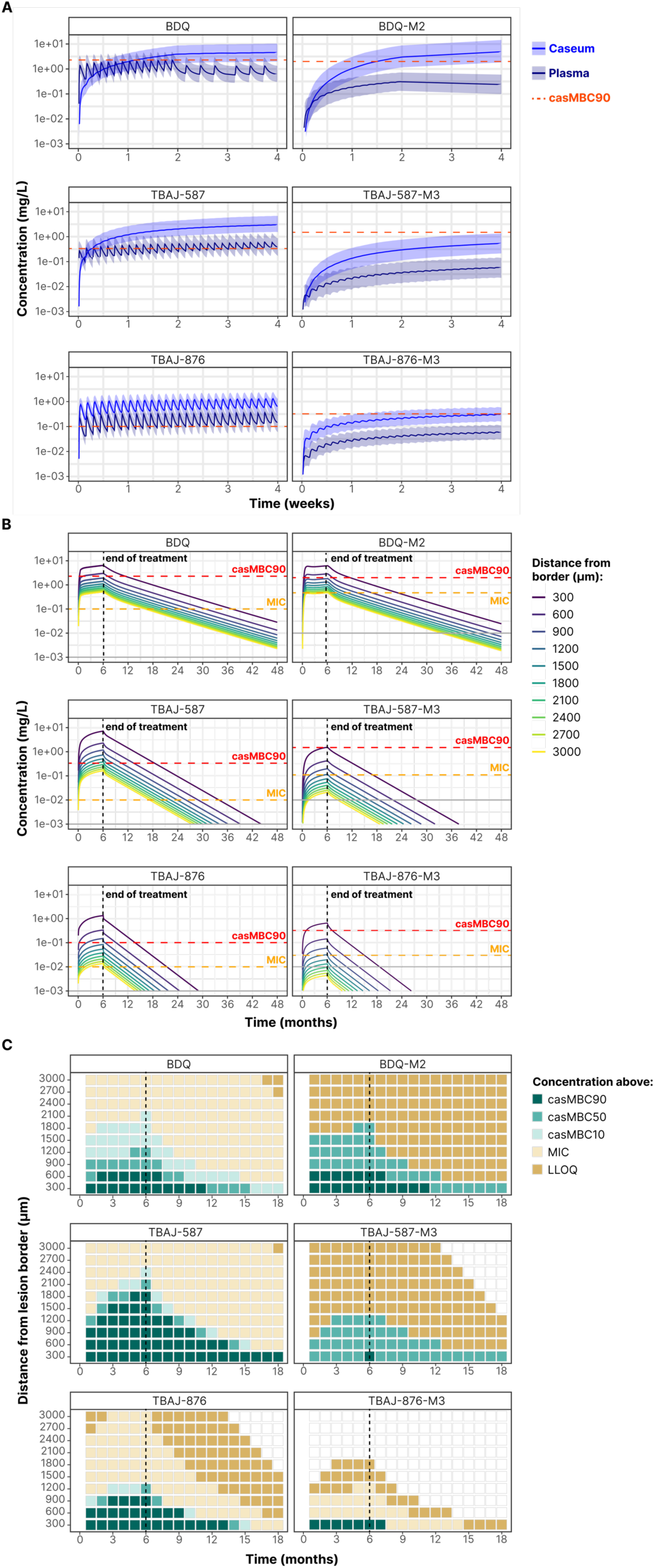
Clinical PK-PD simulations of diarylquinolines and active metabolites in plasma and caseum. Clinical simulations were performed as follows: BDQ at 400 mg QD for two weeks followed by 200 mg thrice weekly for 22 weeks, TBAJ-587 at 200 mg QD and TBAJ-876 at 100 mg QD for 24 weeks. **(A)** Plasma concentrations and lesion exposure in outer caseum (within 300 μm from the cellular rim) were simulated in 500 virtual patients. The shaded area represents the 90% prediction interval of the simulated patient inter-individual variability range, and the solid line is the median patient value. The casMBC_90_ (**Figure S7, Table S2**) is indicated by the red dotted lines. **(B)** Drug exposure as a function of caseum depths in a typical virtual patient during the 24-week treatment and up to 4 years from the start of therapy, relative to the casMBC_90_ and MIC. Gray lines indicate the lower limit of quantitation (LLOQ). **(C)** Target attainment of the diarylquinolines as a function of time from the start of therapy and caseum depth. The dashed black line indicates therapy completion at 6 months. Color coded shaded tiles indicate concentrations achieved at a given time and depth into caseum. Potency targets include casMBC_90_, casMBC_50_, casMBC_10_, MIC, and LLOQ. Blank tiles indicate concentration < LLOQ.

The parent molecules and metabolites reached therapeutic concentrations (casMBC_90_) at the caseum edge (300 μm into the caseous core) at different times from the start of therapy: one week for BDQ, two days for TBAJ-587, one day for TBAJ-876, ten days for metabolite BDQ-M2, and one month for metabolites TBAJ-876-M3. TBAJ-587-M3 did not achieve casMBC_90_ within the first month of therapy (**Figure 3A**). Efficacious exposure at deeper layers into the caseum also varied across the diarylquinolines. TBAJ-587 achieved efficacious concentrations at the greatest depth, followed by TBAJ-876 and BDQ. The TBAJs cleared caseum faster than BDQ after the end of therapy. BDQ and BDQ-M3 persisted at subtherapeutic concentrations in caseum for several years after treatment simulation ended, potentially creating opportunities for resistance development due to de facto monotherapy at sub-bactericidal and ultimately sub-inhibitory concentrations (**Figure 3B**).

Clearance of drug (simulated concentrations below the lower limit of quantitation or LLOQ) from all depths of caseum was not achieved until 72 months after therapy ended for BDQ, while TBAJ-587 and TBAJ-876 cleared within 38 months and 23 months of therapy completion, respectively. BDQ exhibited a gradient of sub-therapeutic exposure deep into the caseum (up to 3000 μm, the maximum simulated depth), which persisted at the deepest level for years after treatment ended. TBAJ-587 exhibited the deepest penetrance (up to 1800 μm) above the casMBC_90_. Among the metabolites, TBAJ-876-M3 and BDQ-M2 had the best exposure above casMBC_90_, and TBAJ-876-M3, like its parent, cleared from all depths of caseum in the shortest timeframe of 12 months post therapy (**Figure 3C**). The faster clearance of TBAJ-876 and TBAJ-876-M3 is likely due to their higher estimated rate of partitioning into caseum (k_lesion_).

Since the TBAJs are approaching treatment shortening Phase 2 trials, we also assessed alternative dosing schemes and whether the differential patterns of diarylquinoline diffusion and clearance would hold true under these scenarios: loading dose (400 mg QD for two weeks followed by 200 mg thrice weekly) versus daily dosing of 200 mg for BDQ, for 4 or 6 months, and varying doses and therapy duration from 2 to 5 months for TBAJ-587 and TBAJ-876. As expected, increasing the TBAJ dose led to deeper PK-PD coverage of caseum at efficacious concentrations. We also observed that the loading dose strategy did not have any significant impact on caseum PK-PD coverage compared to daily dosing (**Figure 4**).

**Figure 4.**
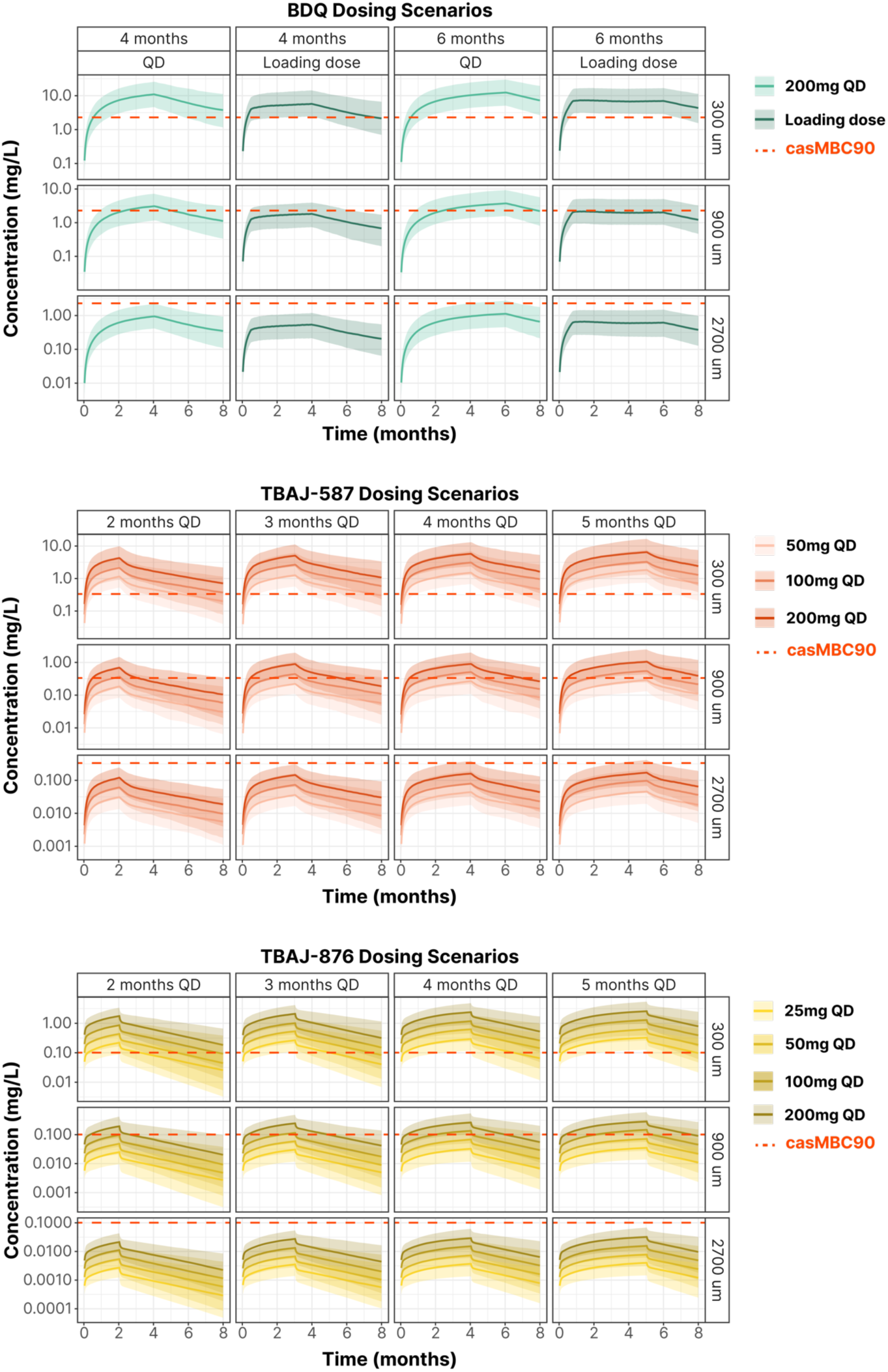
Simulations of diarylquinolines and active metabolites in caseum following clinically relevant dosing regimens. N=500 patients with plasma PK interindividual variability were simulated for each clinically relevant dosing scheme. Shaded areas represent the 90% prediction interval of the simulated patient inter-individual variability range, and the solid line is the median patient value. The three diarylquinolines achieve casMBC_90_ (red dashed line) in outer caseum, and the highest doses of TBAJ-587 (200 mg QD) and TBAJ-876 (100 mg QD) achieve casMBC_90_ at ∼1000 μm. Doses were chosen based on current clinical trials. The loading dose scheme for BDQ is 400 mg QD for two weeks, followed by 200 mg three times weekly for the remainder of treatment (out to four or six months).

## DISCUSSION

Emerging resistance to BDQ threatens its prominent success. Although a historic reservoir of cross-resistance attributed to prior clofazimine exposure is partially responsible (39), resistance acquired during treatment remains the major factor (40). BDQ has a notorious and well described long time to steady state and slow clearance. Here we examined the impact of these PK properties on the kinetics of BDQ diffusion at a crucial non-vascularized site of disease and relapse: caseum or the necrotic center of large lesions.

We found that BDQ achieves steady state and efficacious concentrations in deep caseum after several months and completely clears caseum over several years. This could have positive and negative clinical implications. Under a plausible positive scenario, BDQ and its active metabolite maintain bactericidal concentrations far beyond the end of therapy, potentially eradicating persister cells left behind and preventing relapse. But the slow build up into and clearance out of deep caseum opens two spatio-temporal windows for emergence of resistance, during the first month of therapy and for months to years after concentrations have fallen below the MIC (or MBC against replicating bacteria) but are still sufficient to apply selective pressure and promote expansion of the resistant subpopulation. The active metabolites of BDQ and TBAJ-587 present the largest space-time window of detectable concentrations below their MIC, but that window of space-time is covered by their respective parent (**Figure 3C**), likely neutralizing resistance development. One could argue that acquired resistance in caseum is unlikely since bacterial replication is limited. However, bronchial spread of cavity contents through the airway network (41, 42) exposes this high bacterial burden to varying oxygen tension gradients, conditions favorable to spatio-temporal bursts of replication in a generally non-replicating population. Transient episodes of replication in the context of suboptimal drug concentrations in caseum may be the perfect storm to promote localized emergence of resistance (43). Bronchial spread also shapes connected branched lesions, contributing to disease progression by reseeding the lung (42), and to resistance mutation fixation (44).

That BDQ can linger in caseum for years has multiple causes: a very long half-life in the central compartment associated with a low free fraction in plasma (**Table 3**), an extremely high nonspecific binding in caseum (16) and massive accumulation in foamy macrophages and the cellular rim of necrotic lesions (22), creating a reservoir from which BDQ slowly redistributes into caseum (24). The slow diffusion of BDQ provides an explanation for the disappointing early bactericidal activity after 14 days (25), where efficacy is evaluated in sputum, which partially drains cavity caseum. These site-of-disease PK concepts could similarly apply to other highly hydrophobic TB drugs such as clofazimine.

Modeling the exposure of anti-TB therapeutics at the site-of-action has been conducted in mice and humans (4), including for BDQ, albeit based on a mouse model that does not present necrotic lesions (45). The results presented here are the first to explore target site exposure in caseum – where heavy Mtb burdens are found (9) – as a function of both time and space. The partition coefficient estimated for BDQ at the caseum edge in our model aligns with previously reported BDQ partitioning into whole lung lesions in mice (45). However, our model offers a more comprehensive representation of drug behavior by describing the progressive decline in drug concentration as a function of caseum depth and time. This is afforded by the rabbit model of cavitary TB, which presents a critical feature of human lung pathology (46). Thus, differences in the preclinical species, inherent pathology, experimental methods of tissue sampling and drug quantitation, and modeling strategy preclude direct comparisons of the results.

The design of the present study did not allow for assessing the impact of lesion healing on drug penetration over time, including post-treatment, a difficult question to tackle. While some necrotic foci and lesions heal during treatment, others do not resolve or progress (47, 48), and new ones appear via bronchogenic spread (41, 42). As disease progresses, cavity caseum spills into the airways and is expectorated, leaving a thin caseous layer that adheres to the cavity wall, often connected to the pleural surface. These dirty cavities are the most frequent site of relapse and are located at the apex of the upper and lower lobe where blood flow – and therefore drug supply – is low {Malherbe, 2024 #5254}. This suggests that drug concentrations in outer caseum (∼300-600 μm from the interface with the vascularized cellular rim) may be most relevant to relapse considerations, and that the impact of lower blood flow at the lung apex should be integrated in future models.

Our results indicate that, in comparison to BDQ, the next generation diarylquinolines currently in clinical development reach bactericidal concentrations in superficial and deep caseum more rapidly, and either penetrate deeper layers of caseum (TBAJ-587) or shorten the window of suboptimal concentrations post treatment (TBAJ-876). These desirable properties are expected to reduce opportunities for the emergence of resistance, both in time and space. The simulations of clinically plausible dosing strategies could inform the design of TBAJ-based clinical trials aiming at treatment shortening for patients with cavitary disease.

## METHODS

### Animal infection, drug administration, blood, and lesion sampling

Animal infection studies were performed in Animal Biosafety Level 3 facilities and approved by the Institutional Animal Care and Use Committee of Colorado State University, Fort Collins, CO, the Center for Discovery and Innovation, Hackensack Meridian Health, Nutley, NJ, the National Institute of Allergy and Infection Disease, NIH, Bethesda, MD (LCIM-3E and LCIM-9E). All studies followed the guidelines and basic principles in the United States Public Health Service Policy on Humane Care and Use of Laboratory Animals. All samples collected from *M. tuberculosis* (Mtb) infected animals were handled and processed in the BSL3 in compliance with protocols approved by the Institutional Biosafety Committee of the institutions listed above. C3HeB/FeJ mice, New Zealand White rabbits and common marmosets were infected with Mtb strains Erdman, HN878 and H37Rv, respectively, as previously described (22, 30, 49, 50), until mature lung lesions had developed. Drugs, dosing schedules, and rabbit sampling schemes are described in **Table S1**.

### Lesion sectioning, laser-capture microdissection and drug quantitation

For laser-capture microdissection, large necrotic nodules and cavities were dissected and processed as previously described (51). A laser-capture microdissection scheme was developed to spatially resolve drug quantitation within concentric necrotic areas, each equidistant from the outer border of the caseous core at the interface with the cellular rim (**Figure 1, Figure S8**). The diarylquinolines and metabolites were quantified in plasma and tissue samples by HPLC coupled to tandem mass spectrometry as described in **Supporting Information**.

### Modeling of tissue distribution and PK-PD simulations

For each diarylquinoline, a plasma PK model was built from available infected and uninfected rabbit data. Tissue sample description is provided in **Supporting Information**. Modeling was performed with NONMEM 7.5.0 and PsN 5.3.0. Model building was guided by objective function value, and goodness-of-fit and visual predictive check plots (created using R version 4.2.2 and R packages ggplot2, xpose, xpose4, and vpc). After fixing the final plasma parameters, a site-of-action model was developed for each diarylquinoline where a rate (k_lesion_) and extent of partitioning (PC_lesion_) were estimated for both the parent compound and metabolite (Equation 1).

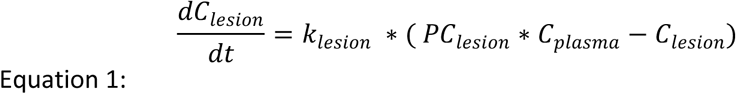

Distance in micrometers from the caseum edge (quantified in LCM experiments described above) was tested as a covariate on k_lesion_ and PC_lesion_ for each diarylquinoline. Multiple covariate relationships were tested including linear, loglinear, power, sigmoidal E_max_, and an inverse relationship. The final model included a distance covariate on the partition coefficient, where DIST is distance in micrometers from the caseum edge (Equation 2).

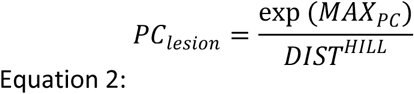

Visual predictive check (VPC) simulations (N=1000) were performed to confirm that the final rabbit lesion models fit the original raw data. Then, to predict drug coverage in patient lesions, the lesion parameters estimated in rabbits were linked to clinical plasma PK models for BDQ, TBAJ-587, and TBAJ-876. The clinical plasma PK models for BDQ and TBAJ-587 were sourced from the literature (36, 37), while TBAJ-876 data (38) were modeled as part of this work. Simulations were compared to potency metrics in a fourteen-day ex vivo caseum assay (casMBC_90_, minimum concentration to kill 90% bacteria (27)), and lesion coverage (drug or metabolite concentration > casMBC_90_) as a function of distance was quantified. Additional casMBC values (casMBC_10_, casMBC_50_, and casMBC_90_) were determined from the fourteen-day ex vivo caseum assay raw data by using a four-parameter logistic model for dose response (dr4pl version 2.0.0). Unless otherwise noted, for all BDQ clinical simulations, virtual patients were given a loading dose of 400 mg BDQ daily for two weeks, followed by 200 mg administered thrice weekly for twenty-two weeks. Unless otherwise noted, simulated patients received 200 mg TBAJ-587 and 100 mg TBAJ-876 daily for twenty-four weeks.

## ACKNOWLEDGEMENTS

This work was supported by shared instrumentation grant S10-OD023524 and S10-OD018072 from NIH to VD, by grant INV-002483 (RS and VD) and INV-040485 (CEB) from the Bill and Melinda Gates Foundation, by grant INV-009105 from the Bill and Melinda Gates Foundation to GTR, and in part by the Division of Intermural Research of the NIAID (LEV and CEB). BDQ was kindly donated by the HIV reagent program of NIH-NIAID and ATCC. BDQ-M2 (N-desmethyl) was provided by Dr Mohamed Nasr, NIH-NIAID. We wish to thank the Global TB Alliance for providing TBAJ-876, TBAJ-587 and their active metabolites. We are grateful to the animal technical teams at Colorado State University and the Center for Discovery and Innovation and the analytical chemistry team of the Center for Discovery and Innovation for their expert technical assistance and to the Tuberculosis Imaging Program personnel of the DIR, NIAID for their technical assistance.

## Notes

### Competing Interest Statement

The authors have declared no competing interest.

