## Supplement for "The kinetics of bedaquiline diffusion in tuberculous cavities opens a window for emergence of resistance"

**This PDF file includes:**

Supporting text
Figures S1 to S8
Tables S1 to S2
SI References

### Supporting Information Text

### METHODS

#### Drug quantitation by HPLC coupled to tandem mass spectrometry

Bedaquiline (BDQ) and BDQ-d6 (Internal Standard, IS) were purchased from Toronto Research Chemicals; verapamil (IS) was purchased from Sigma; BDQ-M2 (N-desmethyl) was provided by Dr Mohamed Nasr, NIH-NIAID. TBAJ876, TBAJ876-M3, TBAJ587 and TBAJ587-M3 were received from the TB Alliance. Drug free $K_2EDTA$ plasma and lung tissue from NZW rabbits and CD-1 mice were obtained from BioIVT for use as blank matrices to build standard curves. For the generation of calibration standards and quality control samples, stock solutions of BDQ, BDQ-M2 and BDQ-d6, TBAJ876, TBAJ876-M3, TBAJ587 and TBAJ587-M3 were prepared in DMSO at a concentration of 1 mg/mL. Working solutions covering the desired concentration range for each drug were prepared by diluting the stock solutions in 50/50 acetonitrile (MeCN)/Milli-Q water (MQW). Drug free plasma or lung homogenate (90 µL) was then spiked with 10 µL of the working solutions to create the calibration standards and quality control (QC) samples. Drugs were extracted from 20 µL of the standard, QC and study samples by the addition of 200 µL of extract solvent containing IS. Extracts were vortexed for 5 minutes and centrifuged at 4,000 rpm for 5 minutes. 100 µL of supernatant was transferred to a 96 well plate for drug quantitation. Processing of LCM samples were prepared by adding 10 µL of each working standard solution to 2 µL of 1:26.7 diluted in PBS drug-free tissue homogenate. Drugs were extracted from the calibration standard, QC, blank/control, and LCM study samples by the addition of 50 µL of extract solvent containing IS. Extracts were sonicated for 10 minutes and centrifuged at 4,000 rpm for 5 minutes. Fifty µL of supernatant were transferred to a 96 well plate for LC-MS/MS analysis. LC-MS/MS analysis was performed on a SCIEX QTRAP 6500+ triple-quadrupole mass spectrometer coupled to a Shimadzu Nexera X2 UHPLC system. Chromatography was performed on an Agilent Zorbax SB-C8 column (2.1x30mm; particle size 3.5 µm) using a reverse phase gradient. MQW deionized water with 0.1% formic acid (FA) was used for the aqueous mobile phase and 0.1 % FA in MeCN for the organic mobile phase. Multiple-reaction monitoring (MRM) of precursor/fragment transitions in electrospray positive-ionization mode was used to quantify the analytes. MRM transitions of 555.00/58.00, 561.00/64.00, 541.00/480.00, 455.40/165.00, 659.00/239.00, 645.00/570.10, 616.00/583.90 and 600.00/525.00 were used for BDQ, BDQ-M2 and BDQ-d6, verapamil, TBAJ876, TBAJ876-M3, TBAJ587, TBAJ587-M3 respectively. Limits of quantification are 1, 10 and 5; 1, 1 and 20; 1, 10 and 2; 1, 5 and 1; 5, 50 and 10; 1, 50 and 50 ng/mL in plasma, tissue and LCM for BDQ, BDQ-M2 and BDQ-d6, TBAJ876, TBAJ876-M3, TBAJ587 and TBAJ587-M3 respectively. Sample analysis was accepted if the concentrations of the quality control samples were within 20% of the nominal concentration. Data processing was performed using the Analyst software (version 1.7.2 Applied Biosystems Sciex).

#### Plasma and tissue samples for model building

For BDQ and BDQ-M2, 55 rabbits contributed 464 plasma measurements, and a subset of rabbits (N=9) contributed 140 lesion data points. Twenty-nine lesion samples (21% of the total lesion samples) were below the limit of quantitation (BLQ), and modeled using the M1 method (1). For TBAJ-587 and TBAJ-587-M3, 23 rabbits contributed 402 plasma measurements, and a subset of rabbits (N=10) contributed 116 lesion data points, respectively. Only two lesion samples (1.7% of the total lesion samples) were BLQ and were thus removed from the analysis (M1 method). For TBAJ-876 and TBAJ-876-M3, 31 rabbits contributed 494 plasma measurements, and a subset of rabbits (N=12) contributed 192 lesion observations. Twenty-seven (14%) of the lesion observations were BLQ, and were again modeled using the M1 Method. The median distance of captured caseum samples from the cellular rim was 440 μm (range 0 to 1796 μm) for BDQ and BDQ-M2, 514 μm (range 78 to 3155 μm) for TBAJ-587 and TBAJ-587-M3, and 545 μm (range 70 to 2527 μm) for TBAJ-876 and TBAJ-876-M3.

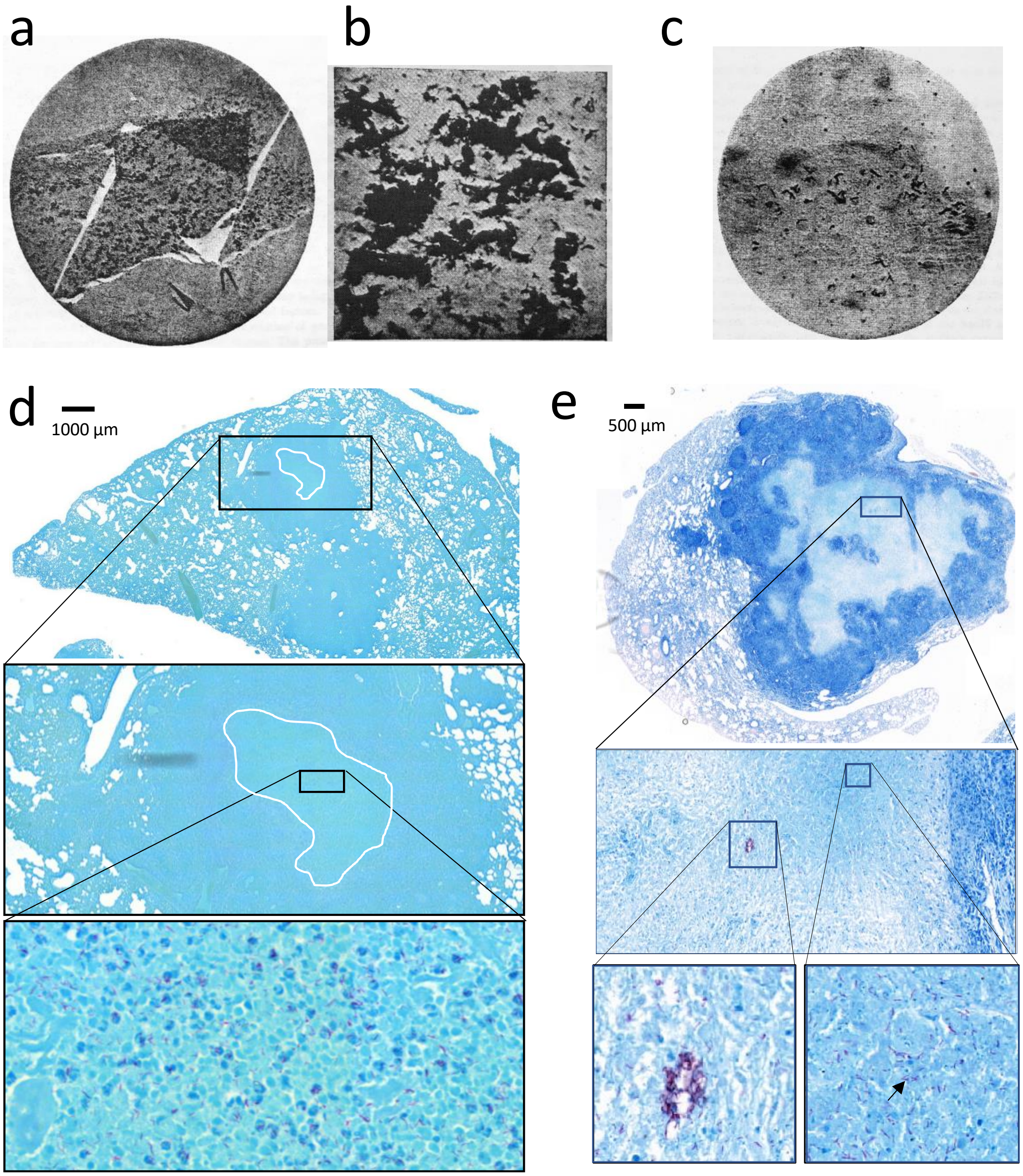


**Fig. S1**. Distribution of *M. tuberculosis* bacilli in the caseum of human and rabbit cavities or large necrotic lesions. **(a)** Mass of bacilli in softening caseum, containing innumerable clumps of bacilli, Ziehl staining x 60. **(b)** Magnification of (a) showing the high density of clusters, Ziehl x 800. **(c)** Necrotic zone at a cavitary surface, showing some clumps. The lumen of the cavity is on the upper right corner, Ziehl x 800.The a-c panels are reproduced from (3). **(d)** Ziehl-Neelsen staining of large necrotic rabbit lesion showing the high density of *M. tuberculosis* bacteria at the center of the caseous mass. **(e)** Ziehl-Neelsen staining of large necrotic rabbit lesion showing a clump reminiscent of the clinical samples shown in (a-c).

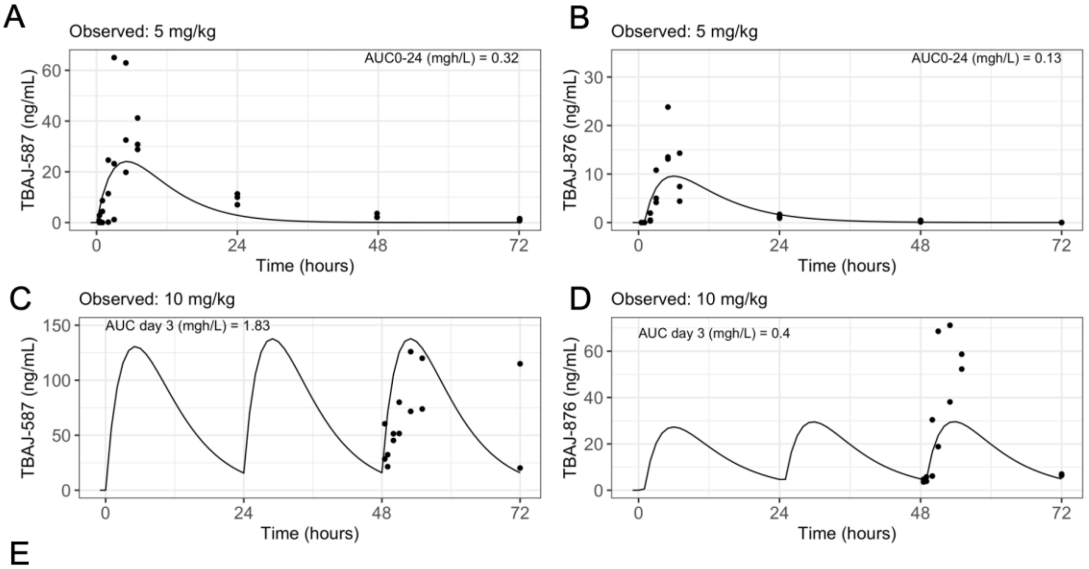


E

| Drug | Species | Dose (mg/kg) | AUC (mg*h/L) |
|---|---|---|---|
| TBAJ-587 | Mouse (target AUC) | 12.5 | 14.8 |
| | Rabbit (observed) | 5 | 0.32 |
| | Rabbit (observed) | 10 | 1.83 |
| | Rabbit (simulated) | 125 | 14.75 |
| TBAJ-876 | Mouse (target AUC) | 3.125 | 4.1 |
| | Rabbit (observed) | 5 | 0.13 |
| | Rabbit (observed) | 10 | 0.40 |
| | Rabbit (simulated) | 100 | 4.13 |

**Fig. S2**. Plasma pharmacokinetics of TBAJ587 and TBAJ876 in uninfected rabbits. In dose finding studies, naïve rabbits received either TBAJ-587 or TBAJ-876, at 5 mg/kg or 10 mg/kg. Based on the observed AUCs, simulations were conducted to determine the dose required to attain a target AUC of 15 μg*h/mL for TBAJ587, and 4 μg*h/mL for TBAJ876, the target efficacious exposure. This resulted in a projected dose of 125 mg/kg of TBAJ-587 and 100 mg/kg of TBAJ-876 for the infected rabbit studies to reach target AUC.

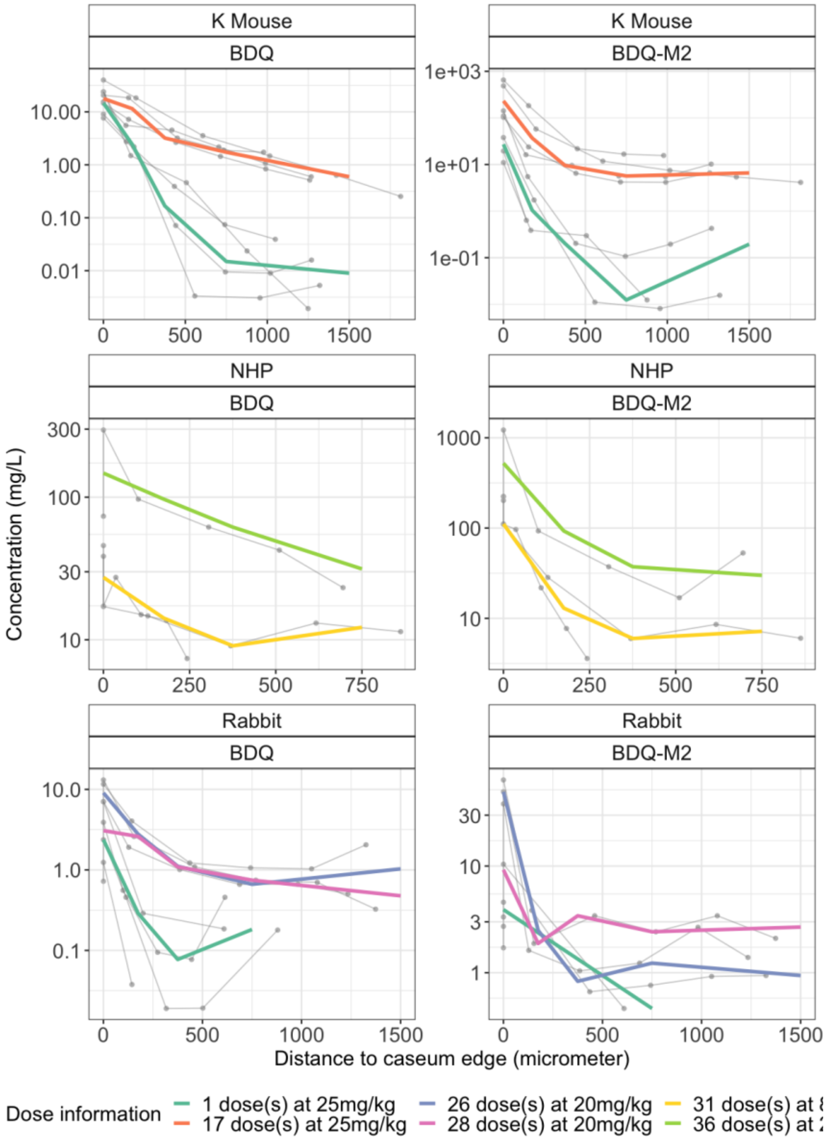


**Fig. S3**. Measurements of diarylquinoline concentrations in laser-capture microdissection (LCM) samples in C3HeB/FeJ mice (K mouse), nonhuman primate (NHP) marmosets and rabbits as a function of caseum depth following various dosing regimens as indicated. Experimental observations are shown as dots, and median of raw data stratified by total number of doses given to the animals. As seen in rabbits, other species also show distant-dependent exposure in caseum, and the steepness of the gradient decreases as the number of doses increases. Mouse and NHP data were not used for subsequent modeling or analysis.

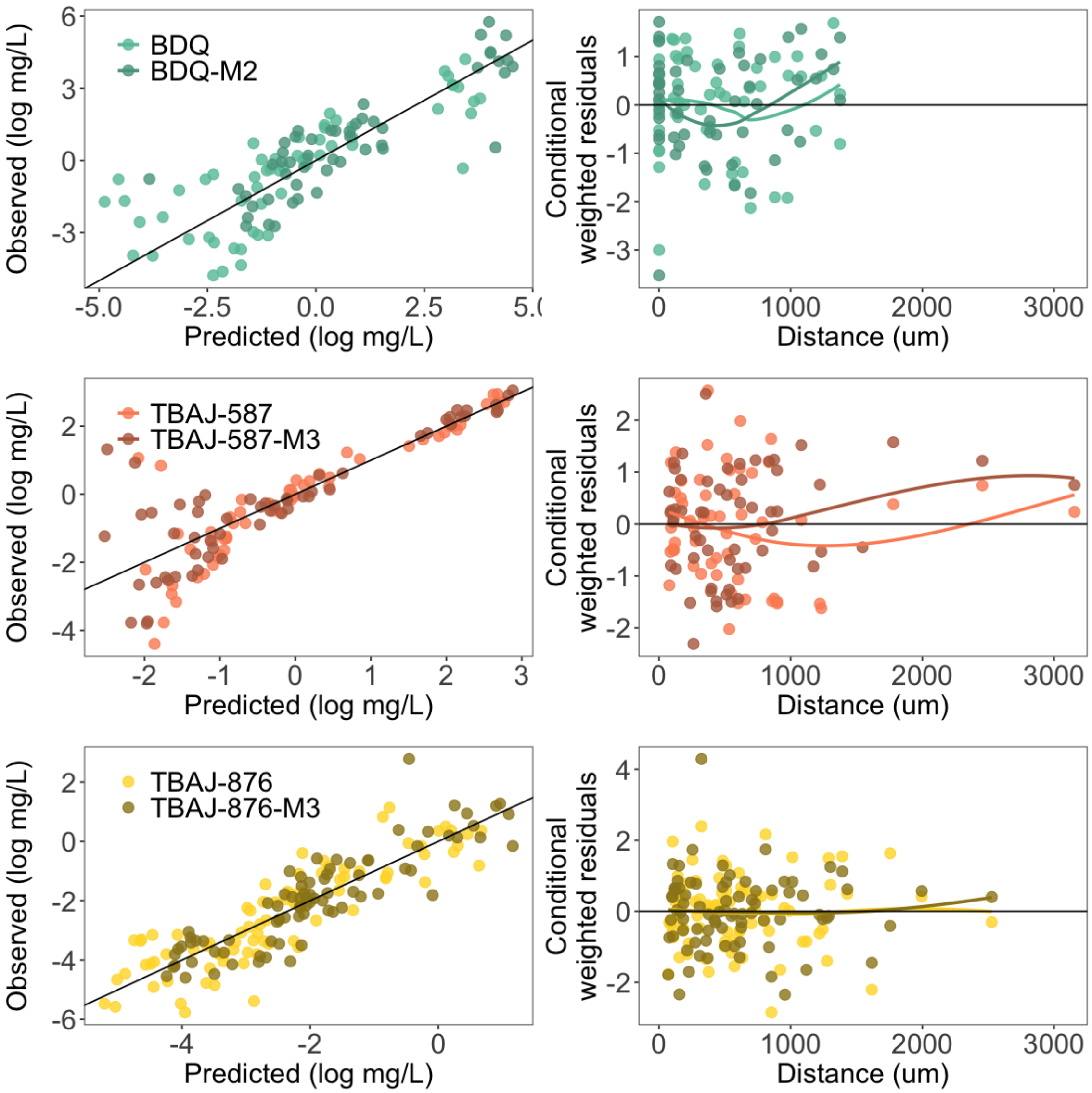


**Fig. S4**. Goodness-of-fit plots for spatiotemporal models. Observed and predicted values were aligned for all diarylquinoline spatiotemporal models. Additionally, the conditional weighted residuals are approximately normally distributed (mean of 0 and a standard deviation of 1), demonstrating that the models are correctly specified.

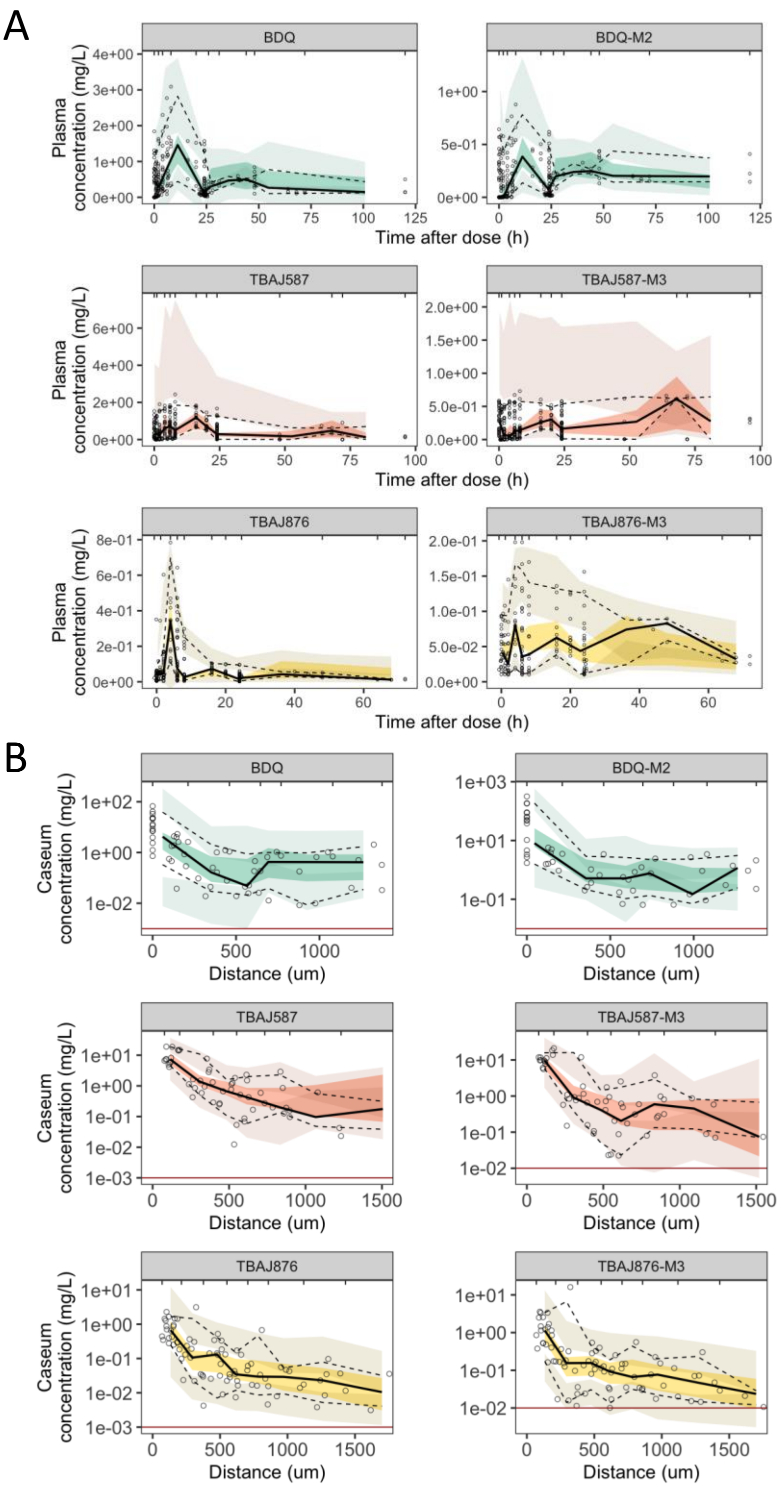


**Fig. S5**. Visual predictive checks of rabbit plasma and lesion models. N=500 simulations of final lesion models were conducted and overlaid with observed data. Observed data is represented as points and median (black, solid line). Simulated data is represented as the 95% confidence interval of the median (dark shaded area). Lightly shaded areas represent the 2.5th and 97.5th percentile of the simulations. The lower limit of quantification is indicated by the red line.

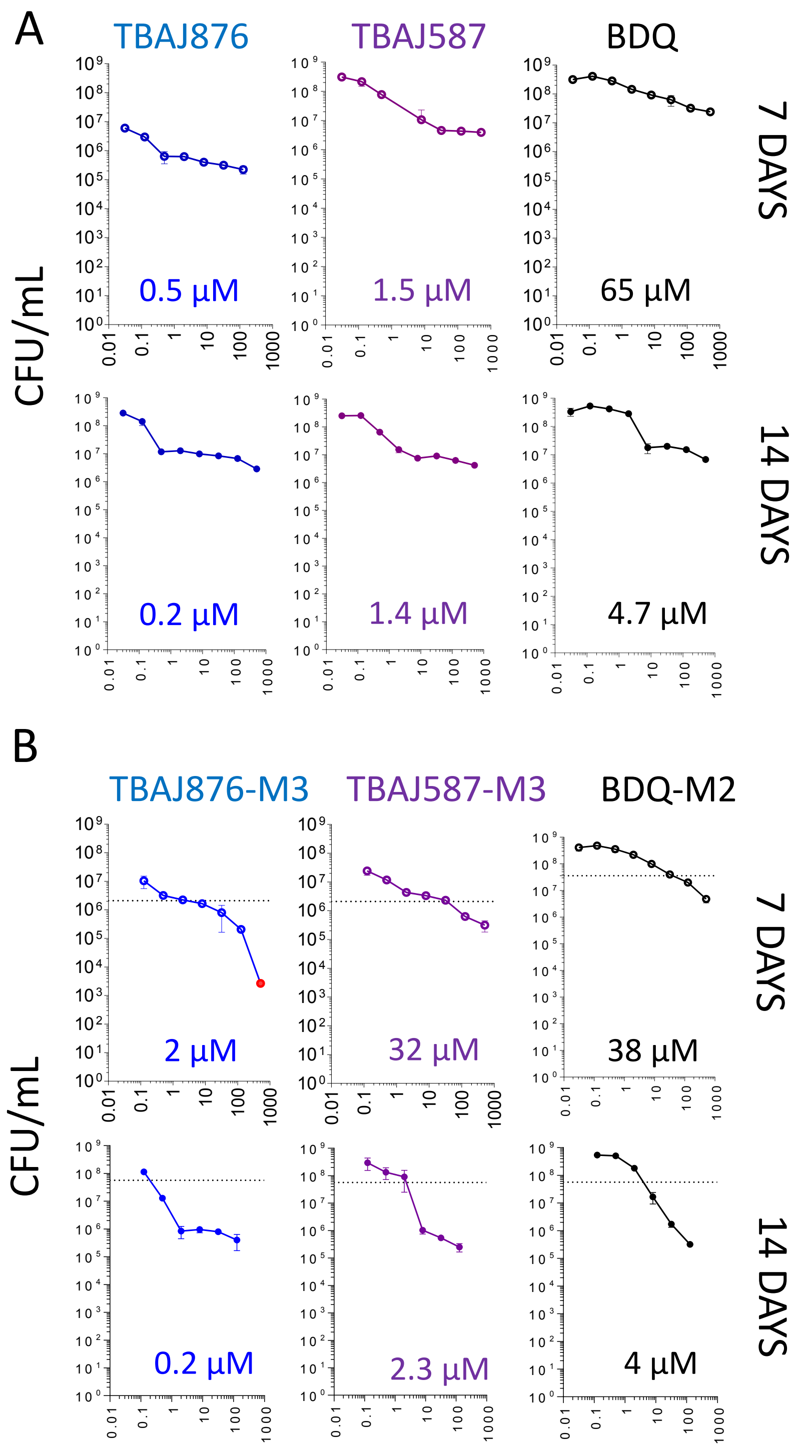


**Fig. S6**. Bactericidal activity of the diarylquinolines and active metabolites against non-replicating *M. tuberculosis* persisters in caseum (4). Due to the slow onset of bedaquiline's bactericidal activity (5, 6), the assay was carried out for 7 and 14 days, showing a marked decrease in casMBC$_{90}$ (estimated from the plots with GraphPad Prism version 10) for bedaquiline and the metabolites. In contrast, the killing activity of TBAJ587 and TBAJ876 is similar after 7 and 14 days.

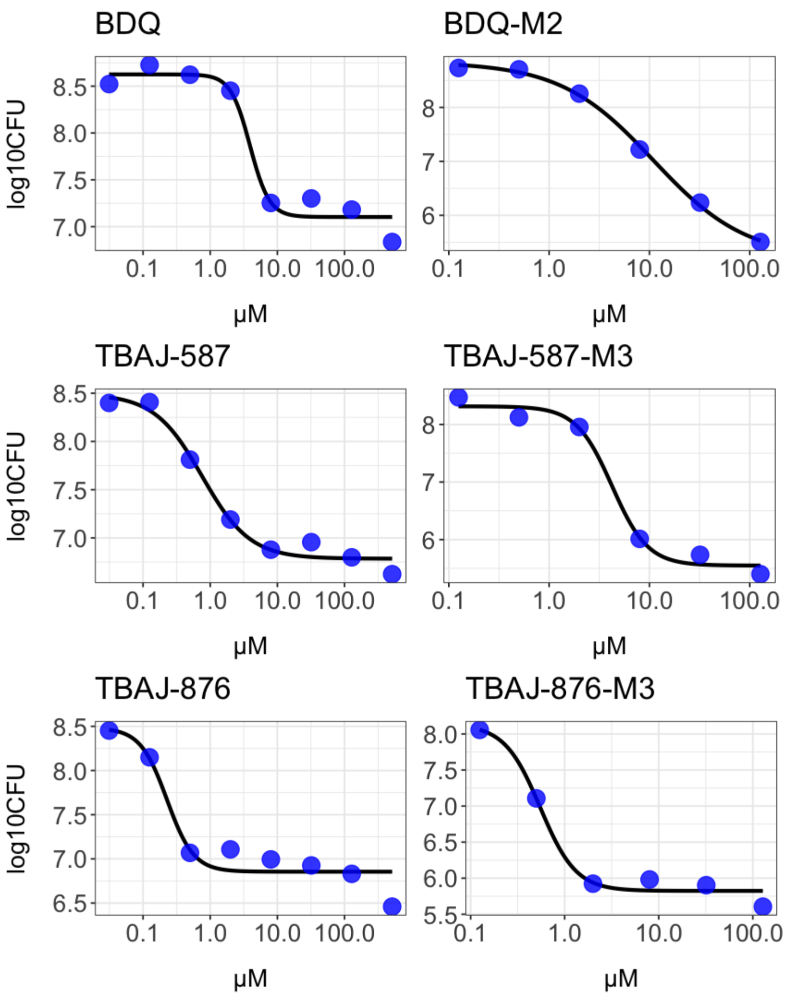


**Fig. S7**. Raw data of 14-day ex vivo caseum assay used as input in the four-parameter logistic dose response model (dr4pl version 2.0.0) to calculate the casMBC$_{90}$, casMBC$_{50}$, and casMBC$_{10}$ (Table S2).

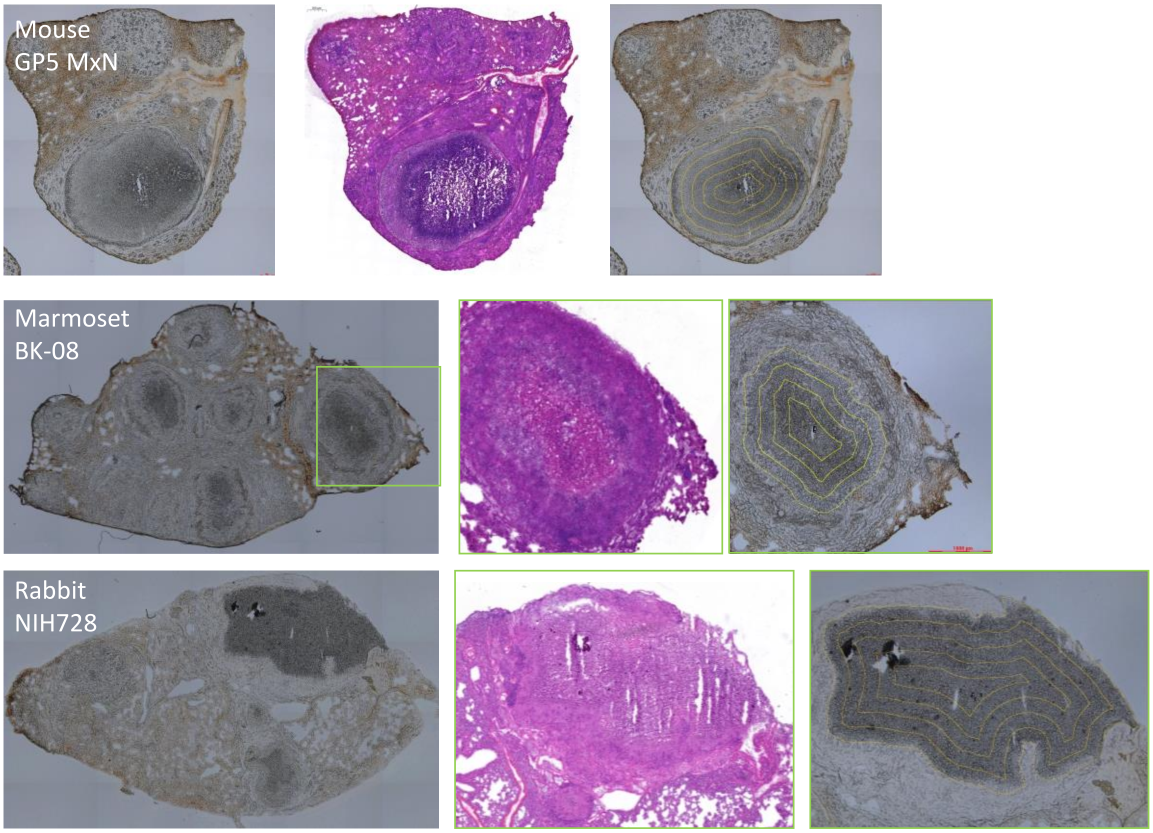


**Fig. S8.** Examples of laser-capture microdissection (LCM) schemes in C3HeB/FeJ mouse (K mouse), nonhuman primate (NHP) marmoset and rabbit samples. Twenty-five μm thick tissue sections were cut from lung lesions using a Leica CM 1860UV cryo-microtome (Buffalo Grove, IL), thaw-mounted onto 1.4 μm thick Leica PET-Membrane FrameSlides (Buffalo Grove, IL), and immediately stored in sealed containers at -80°C. Adjacent 10 μm thick tissue sections were thaw-mounted onto standard glass microscopy slides for hematoxylin and eosin (H&E) histology staining (2). Cellular and necrotic lesion areas were identified optically from the brightfield image scan and by reference to the adjacent H&E image. The Distance Map function of Image J was used to draw equidistant concentric areas on the optical image to guide LCM sampling and calculate the surface of each sampled area. 'Doughnut-shaped' areas totaling 0.5 to $3x10^6$ $μm^2$ were dissected using a Leica LMD7 system (Buffalo Grove, IL). Dissected lesion areas were collected into 0.25 mL standard PCR tubes and immediately transferred to -80°C. The average distance from the outer border of the caseum (in μm, X axis) was calculated by creating a caseum 'mask' and processing it using the Exact Euclidean Distance Transform plugin of the ImageJ software. To determine absolute drug concentrations in each dissected area, the tissue volume was calculated based on the surface area and the 25 μm tissue thickness

**Table S1**. Drugs, dosing schedules, plasma and tissue sampling schemes.

| Rabbit ID | necropsy week | Drug [a] | Dose | # doses | Plasma sampling scheme | TP | Lesions for LCM |
|---|---|---|---|---|---|---|---|
| 168c | 9/21/2020 | BDQ | 400 mg flat dose | 2 | pre, 24, 40, 44, 48h | 48 | 1 |
| 157c | 9/21/2020 | BDQ | 400 mg flat dose | 2 | pre, 24, 40, 44, 48h | 48 | 0 |
| 162c | 9/21/2020 | BDQ | 400 mg flat dose | 2 | pre-dose, 16, 20, 24h | 24 | 0 |
| 164c | 9/21/2020 | BDQ | 400 mg flat dose | 2 | pre-dose, 16, 20, 24h | 24 | 0 |
| 167c | 9/21/2020 | BDQ | 400 mg flat dose | 3 | pre-dose, 16, 20, 24h | 24 | 0 |
| 169c | 9/21/2020 | BDQ | 400 mg flat dose | 3 | pre-dose, 16, 20, 24h | 24 | 6 |
| 171c | 10/12/2020 | BDQ | 400 mg flat dose | 3 | pre, 24, 48, 64, 68, and 72h | 72 | 0 |
| 172c | 10/12/2020 | BDQ | 400 mg flat dose | 3 | pre, 24, 48, 64, 68, and 72h | 72 | 1 |
| 173c | 10/12/2020 | BDQ | 400 mg flat dose | 3 | pre, 0.5, 1, 2, 4, 7h | 7 | 0 |
| 174c | 10/12/2020 | BDQ | 400 mg flat dose | 3 | pre, 0.5, 1, 2, 4, 7h | 7 | 5 |
| 175c | 10/12/2020 | BDQ | 400 mg flat dose | 3 | pre, 24, 40, 44, 48h | 48 | 0 |
| 176c | 10/12/2020 | BDQ | 400 mg flat dose | 3 | pre, 24, 40, 44, 48h | 48 | 3 |
| NIH713 | 8/27/2019 | BDQ | 20 mg/kg | 28 | 24h | 24 | 1 |
| NIH647 | 11/16/2018 | BDQ | 20 mg/kg | 26 | 24h | 24 | 2 |
| NIH723 | 3/25/2020 | BDQ | 20 mg/kg | 1 | 24h | 24 | 1 |
| NIH728 | 3/25/2020 | BDQ | 20 mg/kg | 1 | 24h | 24 | 1 |
| NIH729 | 3/25/2020 | BDQ | 20 mg/kg | 1 | 24h | 24 | 2 |
| 186c | 10/21/2020 | TBAJ876 | 125 mg flat dose | 2 | pre-dose, 16, 20, 24h after 1st and last dose | 24 | 3 |
| 187c | 10/21/2020 | TBAJ876 | 125 mg flat dose | 2 | pre-dose, 16, 20, 24h after 1st and last dose | 24 | 4 |
| 188c | 10/23/2020 | TBAJ876 | 125 mg flat dose | 3 | pre-dose, 16, 20, 24h after 1st and last dose | 24 | 1 |
| 189c | 10/23/2020 | TBAJ876 | 125 mg flat dose | 3 | pre-dose, 16, 20, 24h after 1st and last dose | 24 | 5 |
| 190c | 10/26/2020 | TBAJ876 | 125 mg flat dose | 3 | pre, 16-20-24 after 1st | 72 | 1 |
| 191c | 10/26/2020 | TBAJ876 | 125 mg flat dose | 3 | dose; pre, 24, 48, 64, 68, and 72h after last dose | 72 | 0 |
| 192c | 10/30/2020 | TBAJ876 | 125 mg flat dose | 3 | pre, 4, 6, 8h after 1st | 48 | 0 |
| 193c | 10/30/2020 | TBAJ876 | 125 mg flat dose | 3 | dose; pre, 24, 28, 30, 32 and 48h after last dose | 48 | 0 |
| 194c | 10/28/2020 | TBAJ876 | 125 mg flat dose | 3 | pre, 4, 6, 8h after 1st | 6 | 0 |
| 289R | 11/4/2020 | TBAJ876 | 125 mg flat dose | 3 | dose; pre, 0.5, 1, 2, 4, 6h after last dose | 6 | 1 |
| 290R | 11/13/2020 | TBAJ876 | 125 mg flat dose | 3 | pre-dose, 40, 44, 48 after the last dose | 48 | 2 |

| | | | | | | | |
|---|---|---|---|---|---|---|---|
| 291R | 11/11/2020 | TBAJ876 | 125 mg flat dose | 3 | pre, 4, 6, 8h after 1st dose; pre, 0.5, 1, 2, 4, 6h after last dose | 6 | 0 |
| 302R | 11/12/2020 | TBAJ587 | 300 mg flat dose | 2 | pre-dose, 16, 20, 24h after 1st and last dose | 24 | 0 |
| 303R | 11/12/2020 | TBAJ587 | 300 mg flat dose | 2 | pre-dose, 16, 20, 24h after 1st and last dose | 24 | 0 |
| 304R | 11/19/2020 | TBAJ587 | 300 mg flat dose | 3 | pre-dose, 16, 20, 24h after 1st and last dose | 24 | 0 |
| 305R | 11/19/2020 | TBAJ587 | 300 mg flat dose | 3 | pre-dose, 16, 20, 24h after 1st and last dose | 24 | 2 |
| 306R | 11/16/2020 | TBAJ587 | 300 mg flat dose | 3 | pre, 16-20-24 after 1st dose; pre, 64, 68, and 72h after last dose | 72 | 0 |
| 307R | 11/16/2020 | TBAJ587 | 300 mg flat dose | 3 | | 72 | 2 |
| 308R | 11/20/2020 | TBAJ587 | 300 mg flat dose | 3 | pre, 4, 6, 8h after 1st dose; pre, 2, 4, 6, 8h and 48h after last dose | 48 | 2 |
| 309R | 11/20/2020 | TBAJ587 | 300 mg flat dose | 3 | | 48 | 1 |
| 310R | 11/12/2020 | TBAJ587 | 300 mg flat dose | 3 | pre, 4, 6, 8h after 1st dose; pre, 0.5, 1, 2, 4, 6h after last dose | 6 | 0 |
| 311R | 11/12/2020 | TBAJ587 | 300 mg flat dose | 3 | | 6 | 0 |
| 312R | 11/20/2020 | TBAJ876 | 125 mg flat dose | 3 | pre, 4, 6, 8h after 1st dose; pre, 0.5, 1, 2, 4, 6h after last dose | 6 | 0 |
| 314R | 12/9/2020 | TBAJ876 | 125 mg flat dose | 3 | pre, 4, 6, 8h after 1st dose; pre, 0.5, 1, 2, 4, 6h after last dose | 6 | 3 |
| 317R | 12/9/2020 | TBAJ587 | 300 mg flat dose | 2 | pre-dose, 16, 20, 24h after 1st and last dose | 24 | 0 |
| 318R | 12/16/2020 | TBAJ587 | 300 mg flat dose | 3 | pre, 4, 6, 8h after 1st dose; pre, 0.5, 1, 2, 4, 6h after last dose | 6 | 2 |
| 319R | 12/16/2020 | TBAJ587 | 300 mg flat dose | 3 | | 6 | 0 |
| 320R | 12/9/2020 | TBAJ587 | 300 mg flat dose | 2 | pre-dose, 16, 20, 24h after 1st and last dose | 24 | 1 |
| 248C | 2/19/2021 | TBAJ876 | 125 mg flat dose | 3 | pre-dose, 40, 44, 48 after the last dose | 48 | 0 |
| 249C | 2/18/2021 | TBAJ876 | 125 mg flat dose | 3 | pre, 4, 6, 8h after 1st dose; pre, 0.5, 1, 2, 4, 6h after last dose | 6 | 0 |
| 251C | 2/17/2021 | TBAJ587 | 300 mg flat dose | 2 | pre-dose, 16, 20, 24h after 1st and last dose | 24 | 1 |
| 291C | 9/21/2021 | TBAJ587 | 30.0 mg flat dose | 13 | pre, 2, 6, 8, 24h post last dose | 24 | 1 |
| 292C | 9/21/2021 | TBAJ587 | 30.0 mg flat dose | 13 | pre, 2, 6, 8, 24h post last dose | 24 | 0 |
| 293C | 9/21/2021 | TBAJ587 | 30.0 mg flat dose | 13 | pre, 2, 6, 8, 24h post last dose | 24 | 1 |
| 294C | 10/5/2021 | TBAJ587 | 30.0 mg flat dose | 25 | pre, 2, 6, 8, 24h post last dose | 24 | 0 |
| 295C | 10/5/2021 | TBAJ587 | 30.0 mg flat dose | 25 | pre, 2, 6, 8, 24h post last dose | 24 | 1 |
| 296C | 10/5/2021 | TBAJ587 | 30.0 mg flat dose | 25 | pre, 2, 6, 8, 24h post last dose | 24 | 0 |

| | | | | | | | |
|---|---|---|---|---|---|---|---|
| 297C | 9/22/2021 | TBAJ587 | 30.0 mg flat dose | 13 | pre, 2, 6, 8, 24h post last dose | 24 | 3 |
| 298C | 9/22/2021 | TBAJ587 | 30.0 mg flat dose | 13 | pre, 2, 6, 8, 24h post last dose | 24 | 1 |
| 299C | 9/22/2021 | TBAJ587 | 30.0 mg flat dose | 13 | pre, 2, 6, 8, 24h post last dose | 24 | 0 |
| 300C | 10/6/2021 | TBAJ587 | 30.0 mg flat dose | 25 | pre, 2, 6, 8, 24h post last dose | 24 | 1 |
| 301C | 10/6/2021 | TBAJ587 | 30.0 mg flat dose | 25 | pre, 2, 6, 8, 24h post last dose | 24 | 1 |
| 302C | 10/6/2021 | TBAJ587 | 30.0 mg flat dose | 25 | pre, 2, 6, 8, 24h post last dose | 24 | 3 |
| 303C | 9/23/2021 | TBAJ876 | 12.5 mg flat dose | 13 | pre, 2, 6, 8, 24h post last dose | 24 | 0 |
| 304C | 9/23/2021 | TBAJ876 | 12.5 mg flat dose | 13 | pre, 2, 6, 8, 24h post last dose | 24 | 0 |
| 305C | 10/7/2021 | TBAJ876 | 12.5 mg flat dose | 25 | pre, 2, 6, 8, 24h post last dose | 24 | 0 |
| 306C | 10/7/2021 | TBAJ876 | 12.5 mg flat dose | 25 | pre, 2, 6, 8, 24h post last dose | 24 | 1 |
| 307C | 9/23/2021 | TBAJ876 | 12.5 mg flat dose | 13 | pre, 2, 6, 8, 24h post last dose | 24 | 4 |
| 308C | 10/7/2021 | TBAJ876 | 12.5 mg flat dose | 25 | pre, 2, 6, 8, 24h post last dose | 24 | 0 |
| 309C | 9/24/2021 | TBAJ876 | 12.5 mg flat dose | 13 | pre, 2, 6, 8, 24h post last dose | 24 | 0 |
| 310C | 9/24/2021 | TBAJ876 | 12.5 mg flat dose | 13 | pre, 2, 6, 8, 24h post last dose | 24 | 0 |
| 311C | 10/8/2021 | TBAJ876 | 12.5 mg flat dose | 25 | pre, 2, 6, 8, 24h post last dose | 24 | 0 |
| 312C | 10/8/2021 | TBAJ876 | 12.5 mg flat dose | 25 | pre, 2, 6, 8, 24h post last dose | 24 | 2 |
| 313C | 9/24/2021 | TBAJ876 | 12.5 mg flat dose | 13 | pre, 2, 6, 8, 24h post last dose | 24 | 4 |
| 314C | 10/8/2021 | TBAJ876 | 12.5 mg flat dose | 25 | pre, 2, 6, 8, 24h post last dose | 24 | 0 |
| NIH009 | 10/31/2022 | TBAJ876 | 10 mg/kg | 14 | pre, 1, 2, 6, 24h post last dose | 24 | 5 |
| NIH017 | 10/31/2022 | TBAJ876 | 10 mg/kg | 14 | pre, 1, 2, 6, 24h post last dose | 24 | 8 |
| NIH022 | 10/31/2022 | TBAJ876 | 10 mg/kg | 14 | pre, 1, 2, 6, 24h post last dose | 24 | 4 |
| NIH018 | 11/7/2022 | TBAJ876 | 10 mg/kg | 28 | pre, 1, 2, 6, 24h post last dose | 24 | 8 |
| NIH020 | 11/7/2022 | TBAJ876 | 10 mg/kg | 28 | pre, 1, 2, 6, 24h post last dose | 24 | 4 |

[a] Bedaquiline, TBAJ876 and TBAJ587 were formulated in 20% hydroxypropyl-b-cyclodextrin in 50 mM sodium citrate pH 3 and administered orally at the indicated doses and dosing schedules

**Table S2.** Minimum bactericidal activity of diarylquinolines and metabolites in caseum.

| Compound | Target [a] | Value (μM) [b] |
|---|---|---|
| BDQ | $casMBC_{90}$ | 4.12 |
| BDQ | $casMBC_{50}$ | 2.34 |
| BDQ | $casMBC_{10}$ | 1.32 |
| BDQ-M2 | $casMBC_{90}$ | 3.68 |
| BDQ-M2 | $casMBC_{50}$ | 1.50 |
| BDQ-M2 | $casMBC_{10}$ | 0.61 |
| TBAJ587 | $casMBC_{90}$ | 0.54 |
| TBAJ587 | $casMBC_{50}$ | 0.45 |
| TBAJ587 | $casMBC_{10}$ | 0.37 |
| TBAJ587-M3 | $casMBC_{90}$ | 2.49 |
| TBAJ587-M3 | $casMBC_{50}$ | 0.13 |
| TBAJ587-M3 | $casMBC_{10}$ | 0.01 |
| TBAJ876 | $casMBC_{90}$ | 0.15 |
| TBAJ876 | $casMBC_{50}$ | 0.12 |
| TBAJ876 | $casMBC_{10}$ | 0.10 |
| TBAJ876-M3 | $casMBC_{90}$ | 0.50 |
| TBAJ876-M3 | $casMBC_{50}$ | 0.46 |
| TBAJ876-M3 | $casMBC_{10}$ | 0.42 |

[a] $CasMBC_{90}$, $casMBC_{50}$, and $casMBC_{10}$ values correspond to the minimum concentrations required to kill 90, 50 or 10% of non-replicating bacteria in caseum

[b] Calculated from the 14-day ex vivo caseum assay raw data (**Figure S7 and S8**) using a four-parameter logistic model for dose response (dr4pl version 2.0.0).
